# Learning site-invariant features of connectomes to harmonize complex network measures

**DOI:** 10.1101/2023.09.07.556721

**Authors:** Nancy R. Newlin, Praitayini Kanakaraj, Thomas Li, Kimberly Pechman, Derek Archer, Angela Jefferson, The BIOCARD Study Team, Bennett Landman, Daniel Moyer

## Abstract

Multi-site diffusion MRI data is often acquired on different scanners and with distinct protocols. Differences in hardware and acquisition result in data that contains site dependent information, which confounds connectome analyses aiming to combine such multi-site data. We propose a data-driven solution that isolates site-invariant information whilst maintaining relevant features of the connectome. We construct a latent space that is uncorrelated with the imaging site and highly correlated with patient age and a connectome summary measure. Here, we focus on network modularity. The proposed model is a conditional, variational autoencoder with three additional prediction tasks: one for patient age, and two for modularity trained exclusively on data from each site. This model enables us to 1) isolate site-invariant biological features, learn site context, and 3) re-inject site context and project biological features to desired site domains. We tested these hypotheses by projecting 77 connectomes from two studies and protocols (Vanderbilt Memory and Aging Project (VMAP) and Biomarkers of Cognitive Decline Among Normal Individuals (BIOCARD) to a common site. We find that the resulting dataset of modularity has statistically similar means (p-value <0.05) across sites. In addition, we fit a linear model to the joint dataset and find that positive correlations between age and modularity were preserved.

## 1. INTRODUCTION

Diffusion weighted imaging (DWI) is a non-invasive imaging modality that measures the propensity for water to diffuse in a given direction specified at scan time^1^. With these measurements, we can reconstruct the motions water take in brain tissue. Due to the constraining nature of axons and their bundles (axonal fasciculi), from water motion measurements we can infer the trajectories of these nerve bundles in white matter. Tractography is the process of modelling these trajectories^2,3^. A method of analyzing this representation is called connectomics, which involves the construction of graph reprsentations from tractograms and summarizing the connectivity using complex network measures^4,5^. Complex network measures are used to quantify structural connectivity changes in aging^6^, Alzheimer’s disease^7,8^, epilepsy^9–12^, traumatic brain injuries^13–15^.

There are a growing number of multi-center diffusion imaging studies that span multiple scanner manufacturers and acquisition protocols, or “sites”. Alzheimer’s Disease Neuroimaging Initiative (ADNI)^16^ and National Alzheimer’s Coordinating Center (NACC)^17^ incorporates data from multiple scanner vendors and protocols, Open Access Series of Imaging Studies (OASIS3)^18^ includes multiple protocols, and Baltimore Longitudinal Study of Aging (BLSA)^19^ has data from distinct scanning locations. These site specifications may introduce significant confounding differences in DWI and downstream connectomic analysis^20–23^. Vollmar et. al illustrated confounding site differences in analysis of whole brain, region of interest, and tract-defined microstructure from a traveling subject cohort scanned with the same scanner vendor and protocol^24^. Studies encompassing multiple vendors, models, and protocols had similar findings; these differences result in confounded analyses and are major sources of variation^20^. Further derived metrics such as tractography bundles and complex network measures suffer from these biases as well. Schilling et al. showed that fiber bundle shape and microstructure analysis was affected by scanner manufacturer, acquisition protocol, diffusion sampling scheme, diffusion sensitization and overall bundle processing workflow^22^. Joint datasets of complex network measures in Newlin et al.^23^ and Onicas et al.^25^ show that modularity, global efficiency, clustering coefficient, density, characteristic path length, small worldness, and average betweenness centrality have significant differences due to protocol and scanner vendor. Thus, there is a clear need to account for these site biases in connectivity analyses, or “harmonize”^26–30^.

Previous work on rectifying these biases has explored non-linear harmonization in diffusion MRI, primarily at the image level^31–33^, and more generally across MRI^34,35^. These often rely on removing the predictive information with respect to the site-variable using adversarial losses^33,34^, variational bounds^31^, contrastive losses^36^, and ad hoc disentanglement methods^35^. As shown in^32^, these losses coincide asymptotically as penalties on the mutual information between the learned encoding and the site variable.

We propose to apply a similar approach to removing confounding information from an estimated connectome matrix, via variational bound of the mutual information. The model considers data from multiple sites and learns a representation that is uninformed of those sites. The information we are extracting is a combination of effects due to protocol and study parameters and is therefore not biologically relevant. As such, we force biological information and site information to be disjoint, separable features. With this abstraction, biological connectome information can be projected to any domain by exchanging learned site features. We aim to project data from all sites to one common domain to reduce hardware and acquisition related biases and ultimately harmonize the connectome summary measures. We evaluate the proposed method’s harmonization efficacy by comparing site-wise means and biological variability statistics across the joint dataset.

**Figure 1.**
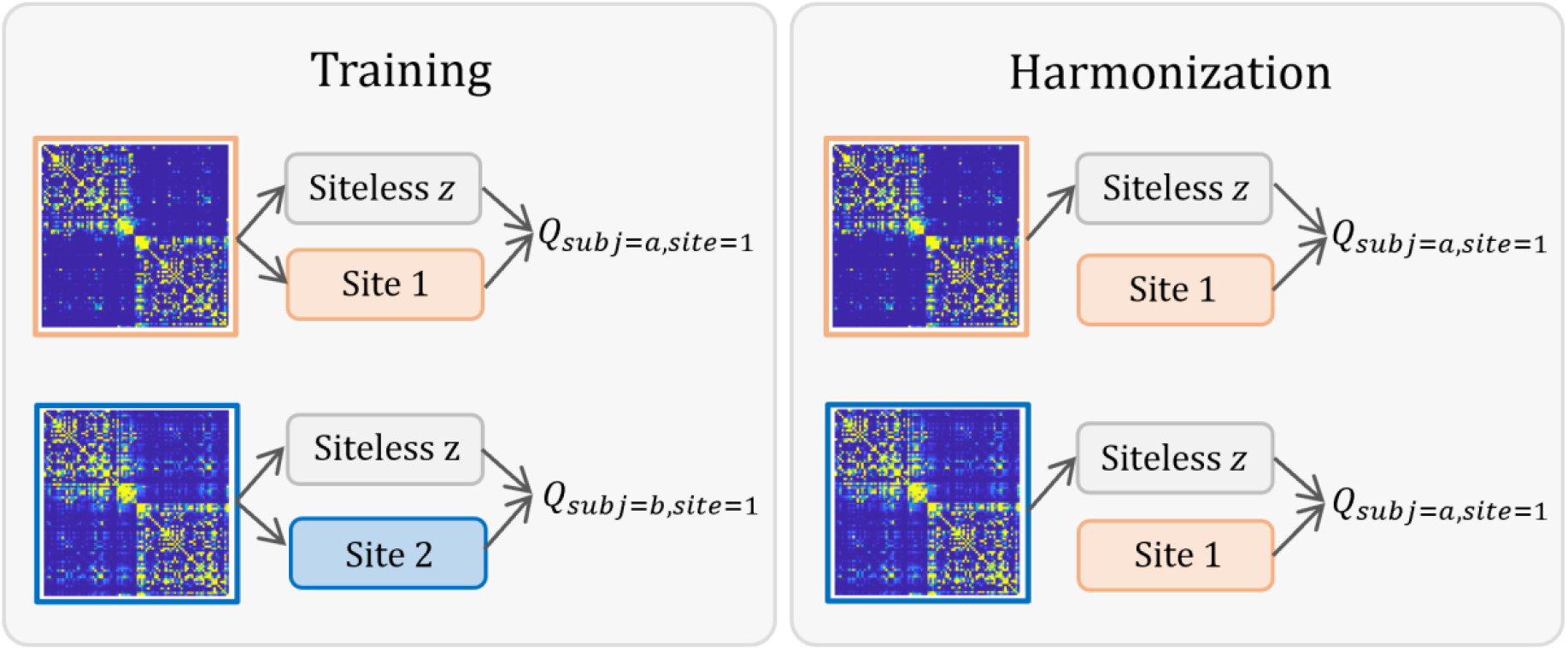
Previous research elucidated that connectomes suffer from confounding site effects. In this work we propose a data-driven model to learn disjoint site (*c*) and biological features (siteless *z*) for BIOCARD (orange) and VMAP (blue) (left). We then inject a prescribed site, *c’*, to the learned representations to compute harmonized connectome modularity, *Q* (right).

## 2. METHODS

We hypothesize that we can disentangle the connectome such that we can extract biological features that have limited or no site information. The site information is a conglomeration of non-biological hardware, protocol, and study parameters that should not drive analysis. We train a neural network that produces representations that are uninformative of the site variable. We then map this representation back to connectivity matrices conditional on a (possibly different) site variable. We show that by manipulating that site variable at test time, we can produce connectivity matrices that at once preserve the biologically relevant information yet also harmonize matrices (i.e., remove site biases). Our method uses multiple latent space prediction heads alongside the main conditional reconstruction task to achieve this site-minimal-bio-maximal representation.

### 2.1. Data

DWI scans from two sites were combined for joint analysis: Vanderbilt Memory and Aging Project (VMAP)^37^ and Biomarkers of Cognitive Decline Among Normal Individuals: the BIOCARD cohort (BIOCARD)^38^. VMAP used a Philips 3T scanner at a resolution of 2 × 2 × 2 mm^3^. BIOCARD used a Philips 3T scanner at a resolution of 0.828 × 0.828 × 2.2 mm^3^.

Data used in this study are split into training and testing cohorts. The training cohort is comprised of subjects 325 scans from VMAP and 692 from BIOCARD, all free of cognitive impairment. Training data from VMAP has ages 74.1 ± 7.5 and 114 women. BIOCARD training data are ages 73.2 ± 5.8 with 410 women. The testing cohort is 77 matched subjects (one scan from each subject), free of cognitive impairment, ages 72.9 ± 7.6, and 57% percent women. The matching was done using pyPheWAS maximal group matching tool (version 4.1.1) ^39^.

### 2.2. Diffusion Processing

The proposed model learns from connectome representations of diffusion tractography and derived from DWI outlined in Section 2.1. DWI from all participants were first preprocessed to remove eddy current, motion, and echo-planar imaging (EPI) distortions prior to any model fitting^40^.

We used the MRTrix default probabilistic tracking algorithm of second order integration over fiber orientation distributions (FODs) for tractography^41^. We generated 10 million streamlines to build each tractogram^42^, limited seeding and termination using the five-tissue-type mask, and allowed backtracking. After, we converted the tractogram to a connectome using the Desikan-Killany atlas^43^ with 84 cortical parcellations from Freesurfer^44^.

We use the Brain Connectivity toolbox (version-2019-03-03) to compute the quality of division of the network into modules, known as modularity^5^. Modularity computed with this toolbox is the site-biased ground truth used in model training.

### 2.3. Model architecture

We implemented a variational-autoencoder (VAE) with site-conditional restrictions on the latent space, z. The overall loss function, *ℓ*_*total*_, is the sum of four sub-component losses: connectome reconstruction (*ℓ*_*recon*_), site-conditional prediction error for modularity for BIOCARD (*ℓ*_*modularity*_(*c*),*c* = 0) and VMAP (*ℓ*_*modularity*_ (*c*),*c* = 1), and age prediction error (*ℓ*_*age*_). Let *x* be the input connectome, 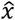 the model reconstruction, *a* the age ground truth, *â* age prediction, *Q* the site-biased modularity and 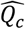 model predicted modularity using layers sequestered for site *c* data.

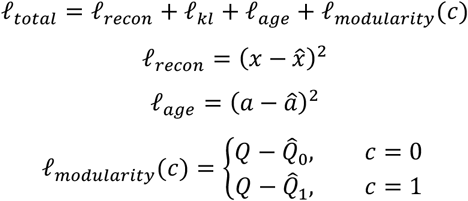

The KL-divergence loss limits site-information in *z*. We would like to remove site-specific information from a learned representation of the connectomes, while remaining maximally relevant with respect to the modularity. Towards that end, our component loss function can be rewritten as mutual information terms (up to constant entropic terms):

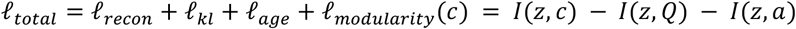

Minimization of the overall implies maximization of each of the negative mutual information terms, which is operationalized as the minimization of predictive error for the second and third term ^45^. Minimization of *I* (*z c*)is achieved using an upper bound which is based on conditional decoding:

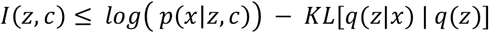

The second term in this bound is difficult to compute directly, but, as shown in Moyer et al. 2020, can be approximated using the standard normal Gaussian in place of induced marginal *q*(*z*)^31^. This fits nicely with exiting VAE literature^46^, is computationally tractable, and, as we show, performs well empirically for removing site information.

**Figure 2.**
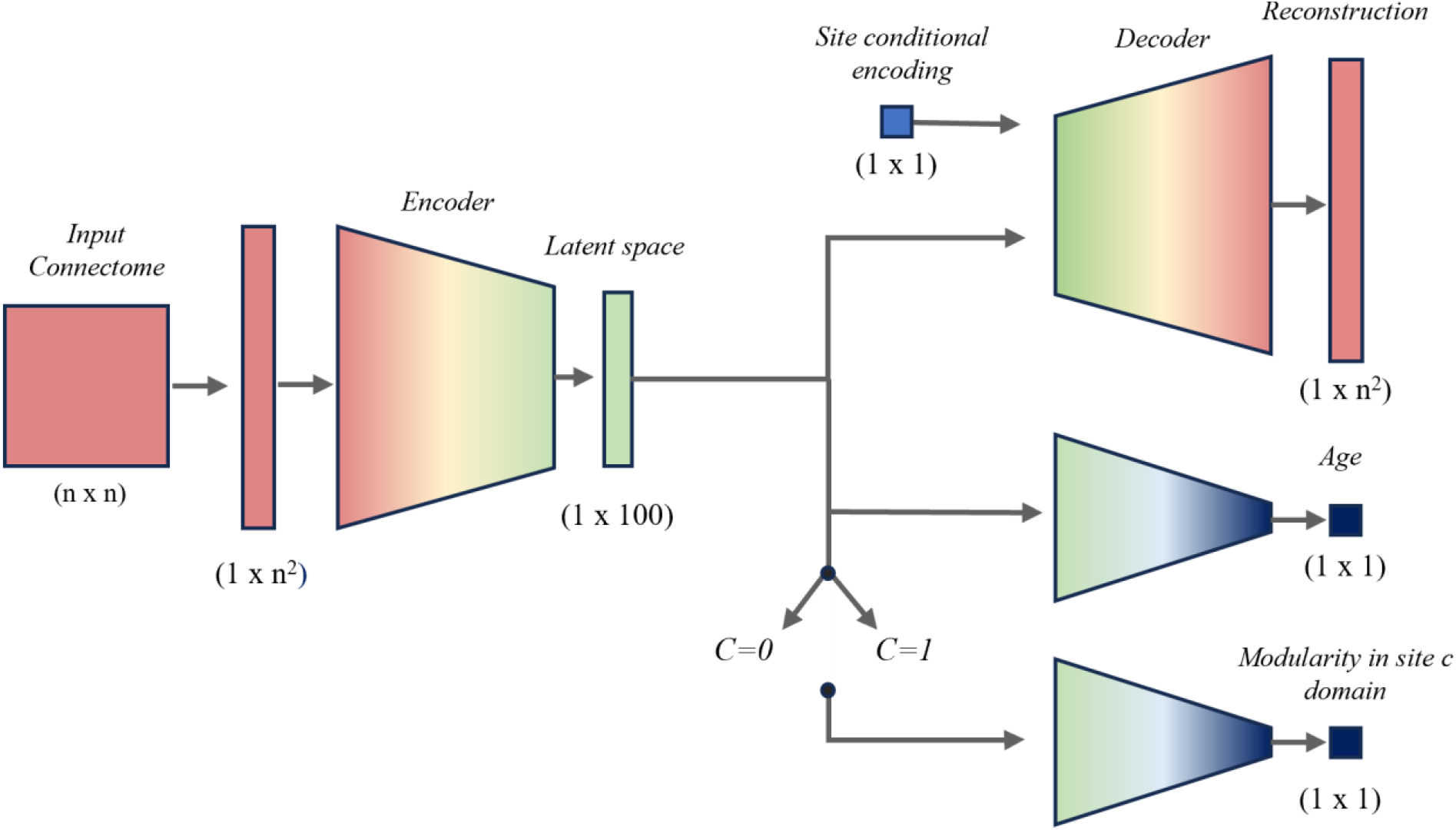
The data are 84 by 84 adjacency matrices weighted by the number of streamlines connecting brain regions corresponding to each matrix edge. This matrix is flattened and passed as input to the model. The reconstruction task has three components: encoding block, latent space, and decoding block. The latent space is partially optimized for reconstruction and used as input for prediction tasks. The prediction tasks are shallow networks that learn calculated, unharmonized modularity of their respective site, *c*. Separate model layers are used for each site. To ground the latent space with biological information, we also predict patient age. The final component contributing to the latent space is kl-divergence loss, which conditions the latent space to have less mutual information with site.

### 2.4. Evaluating harmonization

Harmonization efficacy has two main components: site-invariance and preserved biological information. Site-invariance was evaluated with t-tests for comparing means from each site. We plot modularity against age to assess if biological trends are preserved after multi-site harmonization.

## 3. RESULTS

### 3.1. Site invariance

Using predictive layers trained exclusively on data from one site, we project data from the unseen site onto the desired one. In Figure 3, means are statistically different in A) and after projection to site 1 in B) and site 2 in C), the joint data is harmonized. The data shifts from one site to the other depending on which predictive head is used to generate the modularity values (positive bias for projection to site 1, and negative bias for projection to site 2). We note that variance increases from pre-harmonized to post-harmonized.

**Figure 3.**
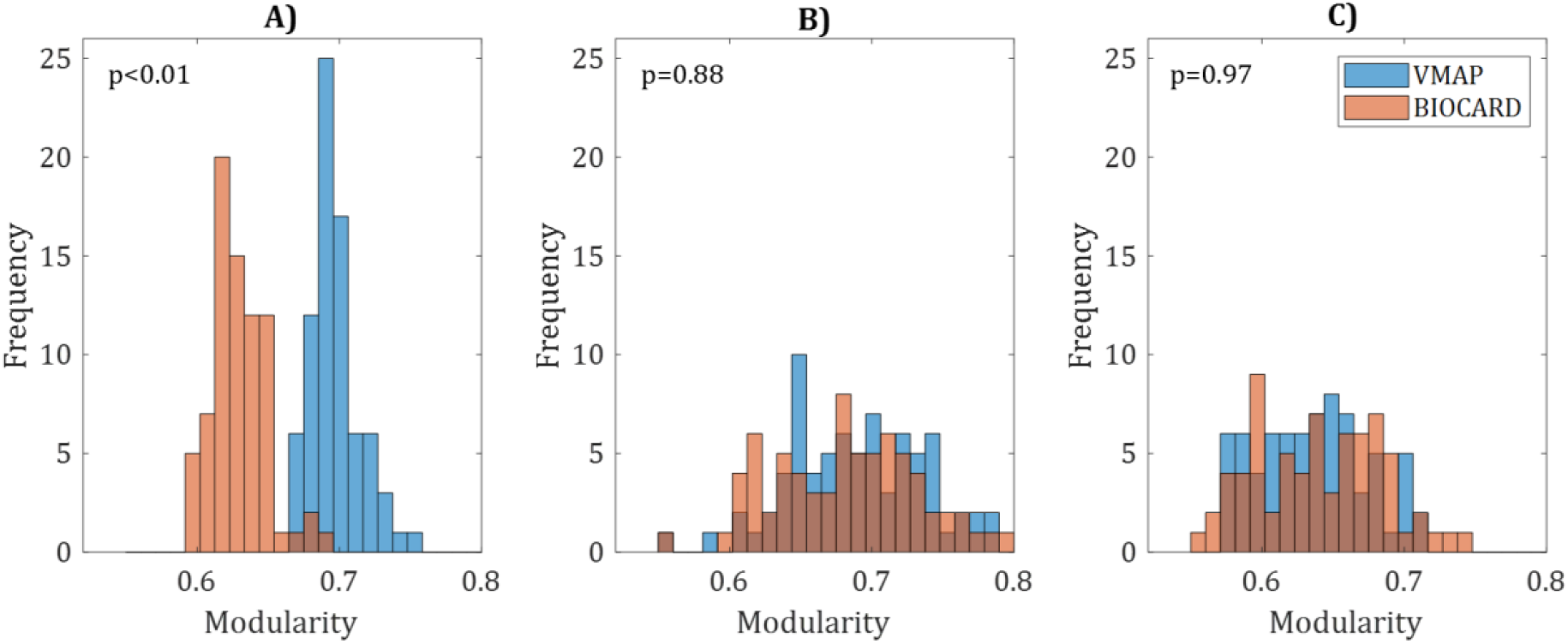
Comparing distributions of modularity from BIOCARD (orange) and VMAP (blue) generated by A) computing modularity using formula on raw connectome generated from tractography, B) predicting from site-invariant latent space and trained on BIOCARD data, and C) predicting from site-invariant latent space and trained on VMAP data. P-values correspond to t-test results comparing means of VMAP and BIOCARD distributions

### 3.2. Preserving biological differences

The top plots in Figure 4 are generated from a single site each (BIOCARD and VMAP, respectively) and therefore do not suffer from any confounding site effects. Their joint, pre-harmonized plot has distinct data clouds with biased intercepts. The data projected to solely VMAP or BIOCARD domain bridge that gap and blend the data together. The harmonization method used to generate such plots preserve the positive correlation (Table 1) with age present in the linear models from each site alone.

**Table 1.** Linear model coefficients with age (*β*)and *R*^2^ for modularity data from each site, pre-harmonized, and predicted from latent spaces trained exclusively on each site. Coefficients have p-values < 0.01.

| Data | Calculated on<br>BIOCARD<br>connectomes | Calculated on<br>VMAP<br>connectomes | Calculated on<br>VMAP and<br>BIOCARD<br>connectomes | Predicted from<br>latent space<br>trained on<br>BIOCARD<br>latent spaces | Predicted from<br>latent space<br>trained on<br>VMAP latent<br>spaces |
| --- | --- | --- | --- | --- | --- |
| $\beta$ | 0.0012 | 0.0015 | 0.0014 | 0.0021 | 0.0031 |
| $R^2$ | 0.294 | 0.37 | 0.0831 | 0.076 | 0.224 |

**Figure 4.**
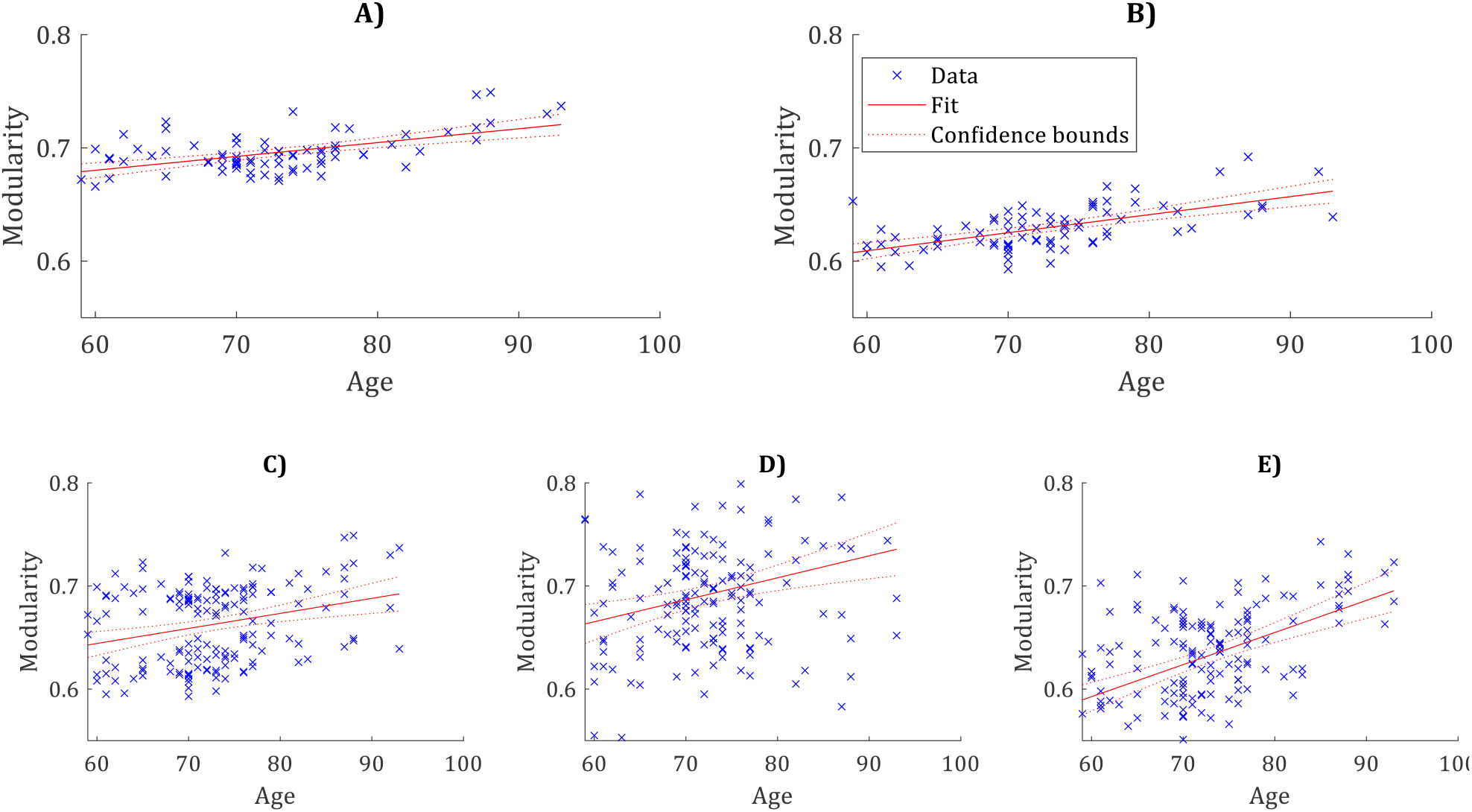
Linear model fit to Modularity A) calculated on VMAP connectomes, B) calculated on BIOCARD connectomes, C) calculated on joint VMAP and BIOCARD connectomes, D) predicted from latent space trained on VMAP latent spaces, and E) predicted from latent space trained on BIOCARD latent spaces.

## 4. CONCLUSION

We discovered that we can 1) use the conditional encoder structure to extract site from the latent space, 2) use that site-less representation and inject new site information, and that 3) we can perform site-injections to produce harmonized modularity. We show it is possible to disentangle site and biological information for the human connectome summary measure, modularity.

## 5. DISCUSSION

Modularity is computed using the connectome and is intrinsically tied to imaging site. To completely remove all site information would remove its semantic value. As such, we are forced to keep at least one site for context if we want to maintain the complex network measure interpretability. Thus, the prediction heads, in being trained exclusively on one site or other, learn the complex domain of the site with respect to modularity. The proposed model enables us to learn and overcome the implicit bias in modularity caused by different imaging protocols. The latent space constructed here can potentially be optimized for other complex network measures as well. However, we hypothesize that not all network measures will be easily disentangled, and some not at all.

## 6. ACKNOWLEDGEMENTS

Data used in preparation of this article were derived from BIOCARD study data, supported by grant U19–AG033655 from the National Institute on Aging. Study data were obtained from the Vanderbilt Memory and Aging Project (VMAP). VMAP data were collected by Vanderbilt Memory and Alzheimer’s Center Investigators at Vanderbilt University Medical Center. This work was supported in part by NIH grant 1R01EB017230, K01-AG073584, NIA grants R01-AG034962, R01-AG056534, and Alzheimer’s Association grant IIRG-08-88733. In addition, this work was supported by Vanderbilt’s High-Performance Computer Cluster for Biomedical Research under award S10-OD023680. The content is solely the responsibility of the authors and does not necessarily represent the official views of the NIH, NIA, or Alzheimer’s Association.

## REFERECNES

[1] Conturo, T. E., Lori, N. F., Cull, T. S., Akbudak, E., Snyder, A. Z., Shimony, J. S., Mckinstry, R. C., Burton, H. and Raichle, M. E., “Tracking neuronal fiber pathways in the living human brain,” Applied Physical Sciences 96, 10422–10427 (1999).

[2] Mori, S. and van Zijl, P. C. M., “Fiber tracking: principles and strategies -a technical review,” NMR Biomed 15(7–8), 468–480 (2002).

[3] Zhang, J., Richards, L. J., Yarowsky, P., Huang, H., van Zijl, P. C. M. and Mori, S., “Three-dimensional anatomical characterization of the developing mouse brain by diffusion tensor microimaging,” Neuroimage 20(3), 1639–1648 (2003).

[4] Bullmore, E. and Sporns, O., “Complex brain networks: graph theoretical analysis of structural and functional systems,” Nat Rev Neurosci 10(3), 186–198 (2009).

[5] Rubinov, M. and Sporns, O., “Complex network measures of brain connectivity: uses and interpretations,” Neuroimage 52(3), 1059–1069 (2010).

[6] Wang, Y., Rheault, F., Schilling, K. G., Beason-Held, L. L., Shafer, A. T., Resnick, S. M. and Landman, B. A., “Longitudinal changes of connectomes and graph theory measures in aging,” Proc SPIE Int Soc Opt Eng 12032, 63 (2022).

[7] Sun, Y., Bi, Q., Wang, X., Hu, X., Li, H., Li, X., Ma, T., Lu, J., Chan, P., Shu, N. and Han, Y., “Prediction of Conversion From Amnestic Mild Cognitive Impairment to Alzheimer’s Disease Based on the Brain Structural Connectome,” article, Front Neurol 9, 1178–1178 (2019).

[8] Ebadi, A., Dalboni da Rocha, J. L., Nagaraju, D. B., Tovar-Moll, F., Bramati, I., Coutinho, G., Sitaram, R. and Rashidi, P., “Ensemble classification of Alzheimer’s disease and mild cognitive impairment based on complex graph measures from diffusion tensor images,” Front Neurosci 11(FEB) (2017).

[9] DeSalvo, M. N., Douw, L., Tanaka, N., Reinsberger, C. and Stufflebeam, S. M., “Altered structural connectome in temporal lobe epilepsy,” Radiology 270(3), 842–848 (2014).

[10] Gleichgerrcht, E., Kocher, M. and Bonilha, L., “Connectomics and graph theory analyses: Novel insights into network abnormalities in epilepsy,” Epilepsia 56(11), 1660–1668 (2015).

[11] Bernhardt, B. C., Chen, Z., He, Y., Evans, A. C. and Bernasconi, N., “Graph-Theoretical Analysis Reveals Disrupted Small-World Organization of Cortical Thickness Correlation Networks in Temporal Lobe Epilepsy,” Cerebral Cortex 21(9), 2147–2157 (2011).

[12] Ibrahim, G. M., Morgan, B. R., Lee, W., Smith, M. Lou Donner, E. J., Wang, F., Beers, C. A., Federico, P., Taylor, M. J., Doesburg, S. M., Rutka, J. T. and Carter Snead III, O., “Impaired development of intrinsic connectivity networks in children with medically intractable localization-related epilepsy,” article, Hum Brain Mapp 35(11), 5686–5700 (2014).

[13] Yuan, W., Wade, S. L. and Babcock, L., “Structural Connectivity Abnormality in Children with Acute Mild Traumatic Brain Injury using Graph Theoretical Analysis” (2014).

[14] Imms, P., Clemente, A., Cook, M., D’Souza, W., Wilson, P. H., Jones, D. K. and Caeyenberghs, K., “The structural connectome in traumatic brain injury: A meta-analysis of graph metrics,” article, Neurosci Biobehav Rev 99, 128–137 (2019).

[15] Caeyenberghs, K., Leemans, A., de Decker, C., Heitger, M., Drijkoningen, D., vander Linden, C., Sunaert, S. and Swinnen, S. P., “Brain connectivity and postural control in young traumatic brain injury patients: A diffusion MRI based network analysis,” Neuroimage Clin 1(1), 106 (2012).

[16] “ADNI | ADNI 3.”, <https://adni.loni.usc.edu/adni-3/> (29 August 2023).

[17] “National Alzheimer’s Coordinating Center.”, <https://naccdata.org/> (29 August 2023).

[18] Marcus, D. S., Wang, T. H., Parker, J., Csernansky, J. G., Morris, J. C. and Buckner, R. L., “Open Access Series of Imaging Studies (OASIS): Cross-sectional MRI data in young, middle aged, nondemented, and demented older adults,” J Cogn Neurosci 19(9), 1498–1507 (2007).

[19] Ferrucci, L., “The baltimore longitudinal study of aging (BLSA): A 50-year-long journey and plans for the future,” Journals of Gerontology -Series A Biological Sciences and Medical Sciences 63(12), 1416–1419 (2008).

[20] Magnotta, V. A., Matsui, J. T., Liu, D., Johnson, H. J., Long, J. D., Bradley D. Bolster, Jr., Mueller, B. A., Lim, K., Mori, S., Helmer, K. G., Turner, J. A., Reading, S., Lowe, M. J., Aylward, E., Flashman, L. A., Bonett, G. and Paulsen, J. S., “MultiCenter Reliability of Diffusion Tensor Imaging,” Brain Connect 2(6), 345 (2012).

[21] Vollmar, C., O’Muircheartaigh, J., Barker, G. J., Symms, M. R., Thompson, P., Kumari, V., Duncan, J. S., Richardson, M. P. and Koepp, M. J., “Identical, but not the same: Intra-site and inter-site reproducibility of fractional anisotropy measures on two 3.0 T scanners,” Neuroimage 51(4), 1384–1394 (2010).

[22] Schilling, K. G., Tax, C. M. W., Rheault, F., Hansen, C., Yang, Q., Yeh, F. C., Cai, L., Anderson, A. W. and Landman, B. A., “Fiber tractography bundle segmentation depends on scanner effects, vendor effects, acquisition resolution, diffusion sampling scheme, diffusion sensitization, and bundle segmentation workflow,” Neuroimage 242, 118451 (2021).

[23] Newlin, N. R., Cai, L. Y., Yao, T., Archer, D., Schilling, K. G., Hohman, T. J., Pechman, K. R., Jefferson, A., Shafer, A. T., Resnick, S. M. and Landman, B. A., “Comparing voxel- and feature-wise harmonization of complex graph measures from multiple sites for structural brain network investigation of aging,” 10.1117/12.265394712464, I. Išgum and O. Colliot, Eds., 537–543 (2023).

[24] Vollmar, C., O’Muircheartaigh, J., Barker, G. J., Symms, M. R., Thompson, P., Kumari, V., Duncan, J. S., Richardson, M. P. and Koepp, M. J., “Identical, but not the same: intra-site and inter-site reproducibility of fractional anisotropy measures on two 3.0T scanners,” Neuroimage 51(4), 1384–1394 (2010).

[25] Onicas, A. I., Ware, A. L., Harris, A. D., Beauchamp, M. H., Beaulieu, C., Craig, W., Doan, Q., Freedman, S. B., Goodyear, B. G., Zemek, R., Yeates, K. O. and Lebel, C., “Multisite Harmonization of Structural DTI Networks in Children: An A-CAP Study,” Front Neurol 13 (2022).

[26] Fortin, J. P., Parker, D., Tunç, B., Watanabe, T., Elliott, M. A., Ruparel, K., Roalf, D. R., Satterthwaite, T. D., Gur, R. C., Gur, R. E., Schultz, R. T., Verma, R. and Shinohara, R. T., “Harmonization of multisite diffusion tensor imaging data,” Neuroimage 161, 149–170 (2017).

[27] Pinto, M. S., Paolella, R., Billiet, T., Van Dyck, P., Guns, P. J., Jeurissen, B., Ribbens, A., den Dekker, J. and Sijbers, J., “Harmonization of Brain Diffusion MRI: Concepts and Methods,” Front Neurosci 14 (2020).

[28] Ning, L., Bonet-Carne, E., Grussu, F., Sepehrband, F., Kaden, E., Veraart, J., Blumberg, S. B., Khoo, C. S., Palombo, M., Kokkinos, I., Alexander, D. C., Coll-Font, J., Scherrer, B., Warfield, S. K., Karayumak, S. C., Rathi, Y., Koppers, S., Weninger, L., Ebert, J., et al., “Cross-scanner and cross-protocol multishell diffusion MRI data harmonization: Algorithms and results,” Neuroimage 221, 117128 (2020).

[29] Mirzaalian, H., Ning, L., Savadjiev, P., Pasternak, O., Bouix, S., Michailovich, O., Grant, G., Marx, C. E., Morey, R. A., Flashman, L. A., George, M. S., McAllister, T. W., Andaluz, N., Shutter, L., Coimbra, R., Zafonte, R. D., Coleman, M. J., Kubicki, M., Westin, C. F., et al., “Inter-site and inter-scanner diffusion MRI data harmonization,” Neuroimage 135, 311–323 (2016).

[30] Tax, C. M., Grussu, F., Kaden, E., Ning, L., Rudrapatna, U., John Evans, C., St-Jean, S., Leemans, A., Koppers, S., Merhof, D., Ghosh, A., Tanno, R., Alexander, D. C., Zappalà, S., Charron, C., Kusmia, S., Linden, D. E., Jones, D. K. and Veraart, J., “Cross-scanner and cross-protocol diffusion MRI data harmonisation: A benchmark database and evaluation of algorithms,” Neuroimage 195, 285–299 (2019).

[31] Moyer, D., Ver Steeg, G., Tax, C. M. W. and Thompson, P. M., “Scanner invariant representations for diffusion MRI harmonization,” Magn Reson Med 84(4), 2174–2189 (2020).

[32] Moyer, D., Gao, S., Brekelmans, R., Steeg, G. Ver and Galstyan, A., “Invariant Representations without Adversarial Training.”

[33] Liu, M., Maiti, P., Thomopoulos, S., Zhu, A., Chai, Y., Kim, H. and Jahanshad, N., “Style Transfer Using Generative Adversarial Networks for Multi-site MRI Harmonization,” Lecture Notes in Computer Science (including subseries Lecture Notes in Artificial Intelligence and Lecture Notes in Bioinformatics) 12903 LNCS, 313–322 (2021).

[34] Kamnitsas, K., Baumgartner, C., Ledig, C., Newcombe, V., Simpson, J., Kane, A., Menon, D., Nori, A., Criminisi, A., Rueckert, D. and Glocker, B., “Unsupervised domain adaptation in brain lesion segmentation with adversarial networks,” Lecture Notes in Computer Science (including subseries Lecture Notes in Artificial Intelligence and Lecture Notes in Bioinformatics) 10265 LNCS, 597–609 (2016).

[35] Zuo, L., Dewey, B. E., Liu, Y., He, Y., Newsome, S. D., Mowry, E. M., Resnick, S. M., Prince, J. L. and Carass, A., “Unsupervised MR harmonization by learning disentangled representations using information bottleneck theory,” Neuroimage 243, 118569 (2021).

[36] Nath, V., Parvathaneni, P., Hansen, C. B., Hainline, A. E., Bermudez, C., Remedios, S., Blaber, J. A., Schilling, K. G., Lyu, I., Janve, V., Gao, Y., Stepniewska, I., Rogers, B. P., Newton, A. T., Davis, L. T., Luci, J., Anderson, A. W. and Landman, B. A., “Inter-Scanner Harmonization of High Angular Resolution DW-MRI using Null Space Deep Learning,” Computational diffusion MRI : MICCAI Workshop 2019, 193 (2019).

[37] Jefferson, A. L., Gifford, K. A., Acosta, L. M. Y., Bell, S. P., Donahue, M. J., Davis, L. T., Gottlieb, J., Gupta, D. K., Hohman, T. J., Lane, E. M., Libon, D. J., Mendes, L. A., Niswender, K., Pechman, K. R., Rane, S., Ruberg, F. L., Su, Y. R., Zetterberg, H. and Liu, D., “The Vanderbilt Memory & Aging Project: Study Design and Baseline Cohort Overview,” Journal of Alzheimer’s Disease 52(2), 539–559 (2016).

[38] “BIOCARD Home Page (NS).”, <https://www.biocard-se.org/public/BIOCARD%20Home%20Page.html> (31 July 2023).

[39] Kerley, C. I., Chaganti, S., Nguyen, T. Q., Bermudez, C., Cutting, L. E., Beason-Held, L. L., Lasko, T. and Landman, B. A., “pyPheWAS: A Phenome-Disease Association Tool for Electronic Medical Record Analysis,” Neuroinformatics 20(2), 483–505 (2022).

[40] Cai, L. Y., Yang, Q., Hansen, C. B., Nath, V., Ramadass, K., Johnson, G. W., Conrad, B. N., Boyd, B. D., Begnoche, J. P., Beason-Held, L. L., Shafer, A. T., Resnick, S. M., Taylor, W. D., Price, G. R., Morgan, V. L., Rogers, B. P., Schilling, K. G. and Landman, B. A., “PreQual: An automated pipeline for integrated preprocessing and quality assurance of diffusion weighted MRI images,” Magn Reson Med 86(1), 456–470 (2021).

[41] Tournier, J. D., Smith, R., Raffelt, D., Tabbara, R., Dhollander, T., Pietsch, M., Christiaens, D., Jeurissen, B., Yeh, C. H. and Connelly, A., “MRtrix3: A fast, flexible and open software framework for medical image processing and visualisation,” Neuroimage 202, 116137 (2019).

[42] Newlin, N. R., Rheault, F., Schilling, K. G. and Landman, B. A., “Characterizing Streamline Count Invariant Graph Measures of Structural Connectomes,” Journal of Magnetic Resonance Imaging (2023).

[43] “CorticalParcellation -Free Surfer Wiki.”, <https://surfer.nmr.mgh.harvard.edu/fswiki/CorticalParcellation> (27 June 2022).

[44] Fischl, F. B., “FreeSurfer” (2012).

[45] Tishby, N., Pereira, F. C. and Bialek, W., “The information bottleneck method” (2000).

[46] Kingma, D. P. and Welling, M., “Auto-Encoding Variational Bayes” (2013).

